# Anterior eye development in the brown anole, *Anolis sagrei*

**DOI:** 10.1101/2021.02.15.429783

**Authors:** Ashley M. Rasys, Shana H. Pau, Katherine E. Irwin, Sherry Luo, Douglas B. Menke, James D. Lauderdale

## Abstract

**Background:** Anterior eye development has been explored in different vertebrate species ranging from fish to mammals. However, missing from this diverse group is a representative of reptiles. A promising candidate to fill this void is the brown anole, *Anolis sagrei*, which is easily raised in the laboratory and for which genome editing techniques exist. Here we provide a detailed histological analysis of the development of the anterior structures of the eye in *A. sagrei*, which include the cornea, iris, ciliary body, lens, trabecular meshwork, and sclera ossicles.

**Results:** Development of the anterior segment in Anoles proceeds as for other vertebrates with the lens forming first followed by the cornea, then the iris, ciliary body, trabecular meshwork, and sclera ossicles. The onset of these latter structures occurs first temporally than nasally. Unlike the eyes of mammals and birds, anoles possess a remarkably thin cornea, flat ciliary body, and a trabecular meshwork that lacks an obvious Schlemm’s canal.

**Conclusions:** This study highlights several features present in anoles and represents an important step towards understanding reptile eye development.

**Key Findings:**

- The anole cornea epithelium is thin, composed mainly of a single basal cell layer.
- The ciliary body lacks a ciliary process.
- Iris and ciliary body formation occur in a spatiotemporal fashion, developing first temporally then nasally.
- The anole trabecular meshwork is composed of a spongiform tissue and lacks a Schlemm’s canal.

## Introduction

The anatomical structure of the vertebrate eye has been described in diverse species.^1-3^ With a consistency that gave Darwin pause,^4^ the canonical vertebrate eye is composed of cornea, lens, iris, ciliary body, muscle, and retina encased in sclera. Beginning with Hans Spemann at the start of the twentieth century, decades of experimental work by numerous investigators have revealed much about the mechanisms controlling the formation of structures common to all vertebrate eyes.^5-8^ However, the eye of any given vertebrate is not just an assemblage of common structures. Rather, its construction during development reflects the needs of the user. Consequently, there is rich variation in the structures of the eye between species.

Reptiles are a class of vertebrates with over 10,000 described species and collectively exhibit rich diversity in eye anatomy.^1,9^ Comparative studies between reptiles, birds, mammals and amphibians have the potential to reveal both common features of tetrapod eye development, as well as features that are unique to reptilian species,^10^ but few studies of eye development have been performed in reptiles. This may be due, in part, to challenges in routinely obtaining large numbers of reptile embryos and to the lack of modern genetic and molecular tools that can be used to investigate the genes and pathways involved in reptile eye development.

An emerging system that has recently overcome many of these barriers is the brown anole lizard, *Anolis sagrei*. This lizard has a relatively short developmental period, going from egg-lay to hatching within a month’s time, reproduces weekly and remains reproductively active for several months of the year.^11^ Moreover, genome modification is now possible in these lizards along with the ability to culture embryos for extended periods.^12,13^ Together, these attributes make this lizard an ideal candidate for embryological studies, and we are working to advance the brown anole lizard as a model for investigations of reptilian eye development.

In this, the second of three papers detailing *Anolis* eye development, we focus on the anterior structures of the eye. Although there are a few historical reports about the tissues of the anterior segment of the reptilian eye,^1,9^ the work we report here is, to our knowledge, the first to comprehensively examine the development of the lens, cornea, iris, ciliary body, and associated tissues in a reptile.

## Materials and Methods

### Animals

All experimental procedures involving lizards were conducted in accordance with the National Institutes of Health Guide for the Care and Use of Laboratory Animals under protocols approved and overseen by the University of Georgia Institutional Animal Care and Use Committee. *Anolis sagrei* were maintained in a breeding colony at the University of Georgia following guidelines described by *Sanger et al*., 2008.^14^ Eggs were collected weekly from natural matings and placed in 100 X 15 mm lidded petri dishes containing moist vermiculite and incubated at 28°C and 70% humidity. Both male and female embryos were used for these studies. Adults and hatchlings were euthanized using methods consistent with the American Veterinary Medical Association (AVMA) Guidelines for the Euthanasia of Animals.^15,16^

### Staging

Embryonic development of *Anolis* lizards typically takes place over a 30-33 day period, starting with internal fertilization.^17^ Early embryogenesis proceeds within the oviduct. *A. sagrei* embryos obtained from eggs that were collected after egg-laying were staged as described by *Sanger et al*., 2008.^17^ Embryos younger than those captured by the Sanger staging series were denoted with the prefix “PL” for pre-laying followed by a number. We describe here 5 PL timepoints, which includes the first few embryonic stages of the Sanger staging series (Sanger St 1-3 correspond to PL 3-5). PL stage embryos were collected from gravid adult females following euthanasia.

### Dissection and Histology

Lizard embryos were accessed by opening eggs using iris scissors and blunt forceps in 1x phosphate-buffered saline (PBS; 137 mM NaCl, 2.7 mM KCl, 10 mM Na2HPO4, 1.8 mM KH2PO4, pH 7.4). Staged embryos or eyes were fixed in Bouin’s Fixative at 4°C overnight. Specimens were stored in 70% EtOH at 4 degrees. Lithium carbonate was added to 1x PBS washes and 70% EtOH in Bouin’s fixed tissue to facilitate removal of picric acid. Embryos or isolated eyes were embedded in paraffin, sectioned on a microtome, and stained with hematoxylin and eosin (H & E).^18^ Photomosaic images were generated using a KEYENCE BZ-700 microscope with Keyence image stitching software. Adobe Photoshop CC (2017.01 release) was used to digitally enhance contrast and adjust white balance of images.

## Results

### Anterior structures of the adult eye

All of the anterior structures in the adult eye are visible in horizontal section (axial section in the naso-temporal plane) through the ocular globe, and include the cornea, lens, iris, ciliary body, and structures responsible for draining the aqueous humor from the eye (Figure 1). Externally, the cornea is positioned towards the nasal side of the ocular globe, slightly off the optical axis.^18^ Histologically, the cornea is composed of an outer epithelial layer, middle stromal layer, and inner endothelial layer (Figure 1.1a). The epithelial layer is primarily a simple epithelium composed of cells with thick squamous morphology (Figure 1.1a). This contrasts with the stratified corneal epithelia found in the eyes other reptiles, birds and mammals.^1,19-22^ The stroma can be divided visually into two layers: an anterior layer adjacent to the corneal epithelium, and a posterior layer that is adjacent to the endothelium. The anterior layer, which we believe to be Bowman’s membrane,^9^ exhibits a compact arrangement of collagen fibrils, while the posterior layer has a more relaxed arrangement (Figure 1.1a). Scattered, but typically straddling the midline between these stroma divisions, are presumptive keratocytes. Finally, the endothelial layer of the cornea is composed of a single layer of cells with flattened morphology (Figure 1.1a). We were not able to identify by histology a clearly defined posterior limiting membrane, known as Descemet’s layer, separating the stroma from the endothelium. With the exception of Descemet’s layer, our histological findings are consistent with the written description provided of the cornea for *A. lineatopus*.^9^ As with other reptiles, birds and mammals, the stroma contributes to most of the corneal thickness.^1,19-22^

**Figure 1.**
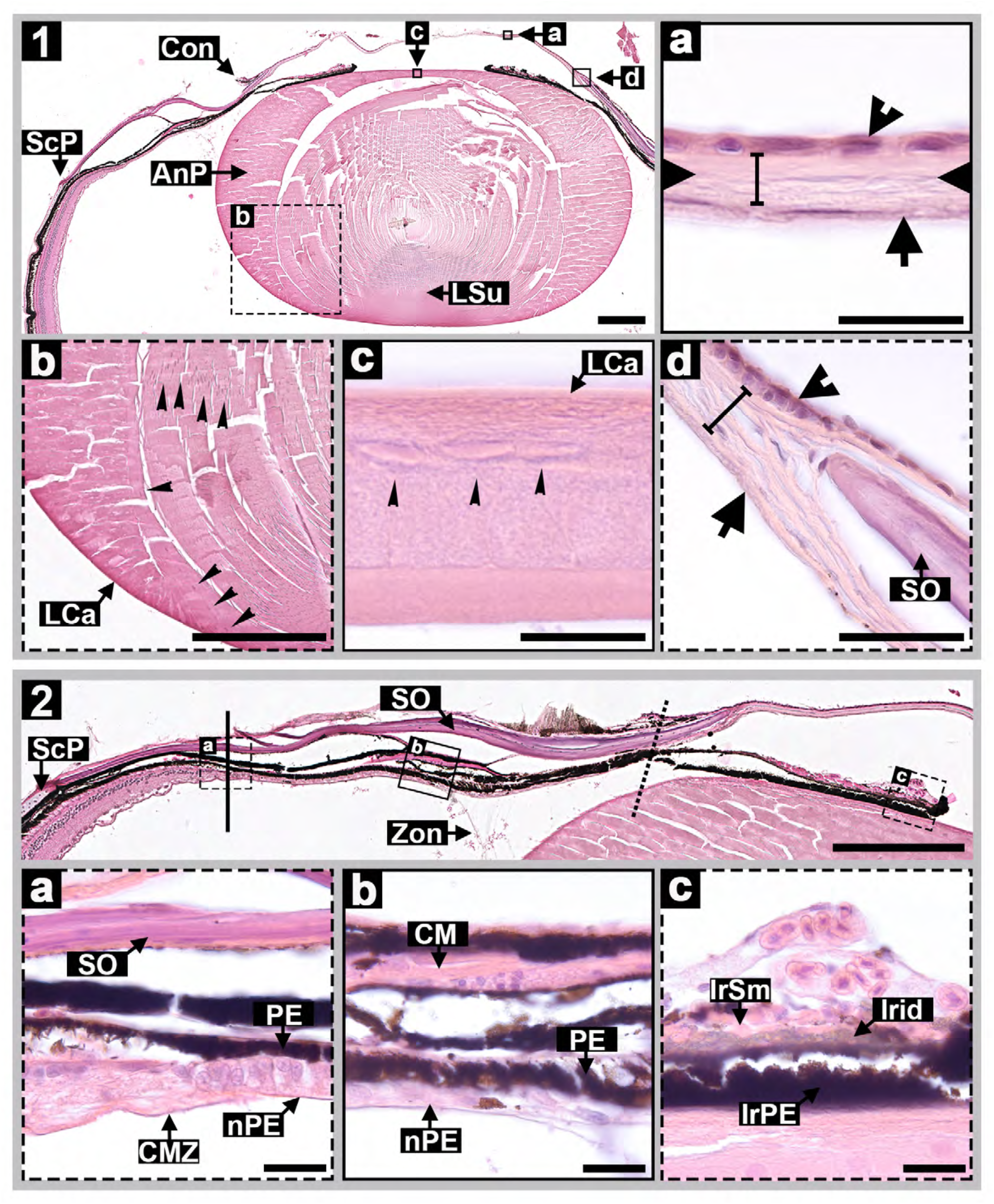
Anterior structures of the adult anole eye. (1) is a horizontal section showing lens, cornea, iris, and ciliary body. (a-d) are high magnification views of cornea (1.a), lens bow (1.b), lens epithelial (1.c), and limbus region (1.d). (1.a-d) markers represent; Con – conjunctiva, ScP – sclera proper, LSu – lens suture, AnP – annular pad, LCa –lens capsule, SO – sclera ossicles, arrowheads – corneal epithelial, bar – stroma, arrow – corneal endothelial layer, notches – bowman’s membrane boarder, and narrow arrow heads – lens epithelial cells. (2) is a horizontal section depicting iris, ciliary body, and ciliary marginal zone; lines indicate ciliary marginal zone (solid) and iris (dashed) boundaries. (2.a-c) are high magnified inserts of the ciliary marginal zone (2.a), ciliary body (2.b), and iris (2.c). Markers; SO – sclera ossicles, Zon – zonular fibers, CMZ – ciliary marginal zone, PE – pigment epithelium, nPE – nonpigmented epithelium, CM – ciliary muscle, IrSm – iris smooth muscle, IrPE – iris pigmented epithelium, and Irid – iridophores. Scale bars 20µm (1.a,c and 2.a-c), 40µm (1.d), and 250µm (1, 1.b, and 2).

The gross structure of the lens of adult *A. sagrei* (Figure 1) is directly comparable to that previously described for *A. lineatopus*.^9^ The lens, which is soft compared to mouse, rabbit, horse, and human (the mammals for which we have lens data), is elliptical in shape with a smaller center nucleus, larger cortex, and outer epithelium (Figure 1.1; Figure 2). The lens nucleus is defined by the presence of primary lens fiber cells, and the cortex is defined by the presence of secondary lens fiber cells. The nucleus and cortex make up the main lens body. The lens epithelium is thicker around the equator where it forms the annular pad, also known as the “Ringwulst” (Figure 1.1; Figure 2). The annular pad is characterized by nucleated fibers and is a feature present in the lenses of many reptiles and birds ^1,9,23,24^. Surrounding these structures is a thick lens capsule (Figure 1.1,b,c). At the anterior face, long thin zonular fibers can be seen anchoring the lens to the ciliary body (Figure 1.2).

**Figure 2.**
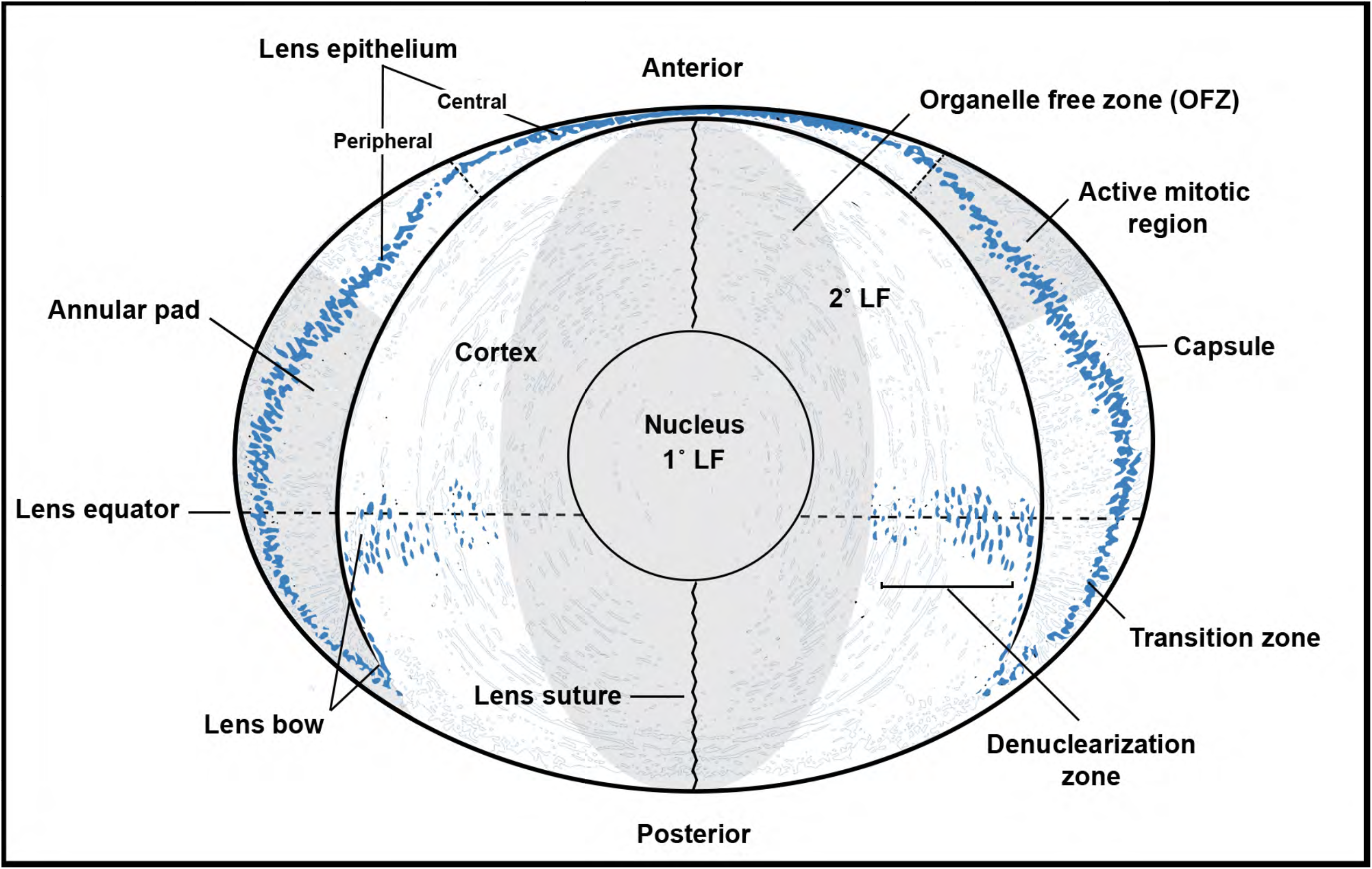
Anatomical diagram of the anole lens. 1°LF – primary lens fiber, 2°LF – secondary lens fiber.

The iris is a muscular structure that controls the amount of light entering the eye. Prominent notches are visible at approximately 12 o’clock and 7 o’clock (right eye) or 12 o’clock and 5 o’clock (left eye) on the iris face when eyes are viewed externally and in their natural position within the skull.^18^ The length of the iris is broader temporally (the region subtended from 12 to 5 o’clock) than nasally (the region from 5 to 12 o’clock).^18^ This asymmetry appears to align the center of the iris aperture with that of the cornea.The iris is closely associated with the lens (Figure 1.2). Histologically, the iris (the pars iridica retinae) is an extension of the ciliary epithelium. It is composed of outer and inner epithelial layers which are thin; individual cell morphology is not easily observed by brightfield microscopy due to the dark pigmentation of these cells. Superficial to these layers is the stroma. The stroma is vascularized, has some pigment granules, and attaches to the sphincter and dilator muscles, which are striated and not smooth (Figure 1.2). Strands of iris/uveal tissue, likely the pectinate ligaments, connect the base of the iris to the peripheral cornea/sclera, forming the boundary of the anterior chamber at the iridocorneal angle.

Located between the iris and ora serrata, the ciliary body is an annular structure located at the equator of the lens. In section, the ciliary body is linear rather than folded like in mammals and birds (Figure 1.2). The ciliary body (the pars ciliaris retinae), like the iris, is comprised of an outer and inner epithelial cell layers. The outer epithelium is heavily pigmented and is contiguous with the pigmented epithelium of the iris on one side and the retinal pigmented epithelium on the other. The inner epithelial layer is non-pigmented. The ciliary muscles, which control the shape of the lens, extend from the sclera and corneal limbal region towards the ciliary body and marginal zone. These muscles are in two groups, one distal and another proximal, represented by Crampton’s and Brücke’s muscles, respectively. Like the iris, these muscles are striated.

In mammals, birds, and presumably reptiles, the ciliary body secretes aqueous humor into the posterior chamber of the eye, which is the space between the iris and lens. The aqueous humor then flows through the pupil to the anterior chamber, which is the space between the iris and cornea. In mammals and birds, the aqueous humor exits the eye via outflow through a porous tissue called the trabecular meshwork, which is located at the angle between the iris and cornea.^25-27^ In mammals, and perhaps birds, the aqueous humor flows from the trabecular meshwork into a vascular structure known as Schlemm’s canal and then into the venous system. ^25-27^ Inspection of the iridocorneal angle in the eyes of adult *A. sagrei*, reveals a spongiform tissue comparable to the trabecular meshwork in chickens.^26^ We were not able to identify a structure comparable to Schlemm’s canal as defined for the mammalian eye.^25^ Instead, we noted that the vasculature from the choroid region extends between the ciliary body and the ciliary muscles to the iridocorneal angle and propose that this contributes to the outflow of aqueous humor.

### Lens development

In anoles, the earliest stages of eye development take place while the eggs are in the oviduct. While it is difficult to provide a precise timeline of development during this period, based on our observations we estimate that the time between fertilization and egg lay is close to 4 days. For comparative purposes, using morphological criteria we have created a staging series that captures embryonic development that occurs in the oviduct. Embryonic stages that occur pre-egg lay are denoted with the prefix “PL”. Stages observed in embryos post-egg lay use the Sanger staging series designations^17^ and are denoted using the prefix ”St”. Stages PL 3-5 correspond to Sanger stages St 1-3. For context, after egg lay it takes another ∼30 days before the lizard hatches.

Embryos at about 1-day post fertilization (PL1), have optic vesicles but no clear evidence of lens placodes.^18^ By PL2, the lens placode is evident in the surface ectoderm overlying the neuroectoderm of the optic vesicle/early optic cup (Figure 3a). By PL3, the lens placode has invaginated to form the lens pit (Figure 3b), which develops into the lens vesicle by PL 4 (Figure 3c). At this time, the lens vesicle appears to be surrounded by a capsule (Figure 3c). At PL5 the lens is becoming polarized as cells in the posterior vesicle elongate and differentiate to form primary lens fiber cells, and cells in the anterior vesicle are differentiating to form the epithelium (Figure 3d). At the time of egg lay (∼St 4), the lens is completely separated from the surface ectoderm and is composed of anterior cuboidal epithelial cells and posterior elongating fiber cells (Figure 3e). The anterior cuboidal epithelial cells have formed a pseudostratified epithelial layer, while the primary fiber cells have completely extended into the vesicle space forming the nucleus of the lens (Figure 4.1,a).

**Figure 3.**
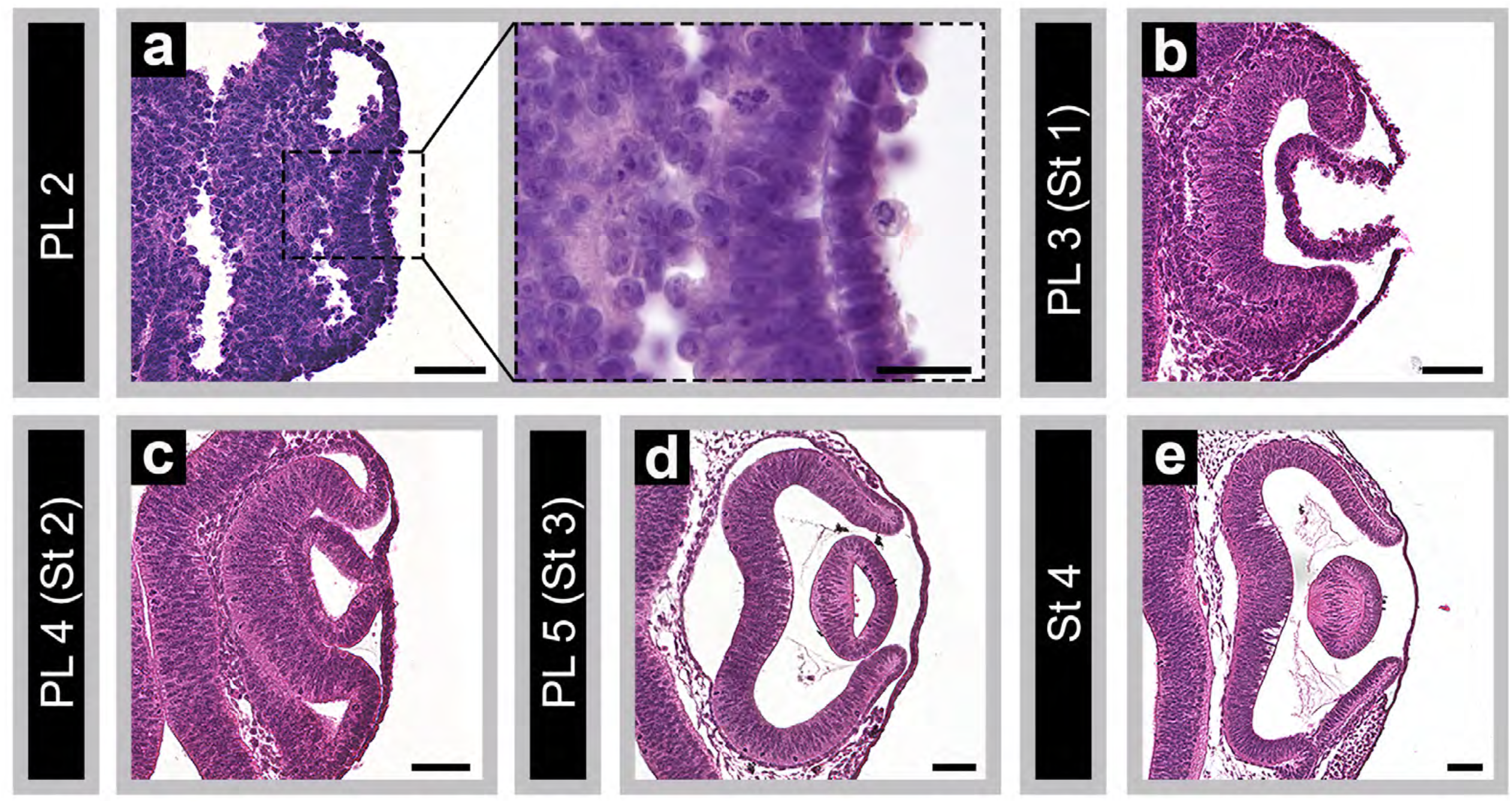
Early anole embryonic eye development. (a-e) are coronal sections from PL2 – Stage 4 embryos depicting lens placode, optic cup (a), lens pit (b), and lens vesicle formation (c-d); Scale bars 50µm (a-e), insert (a) 20µm.

**Figure 4.**
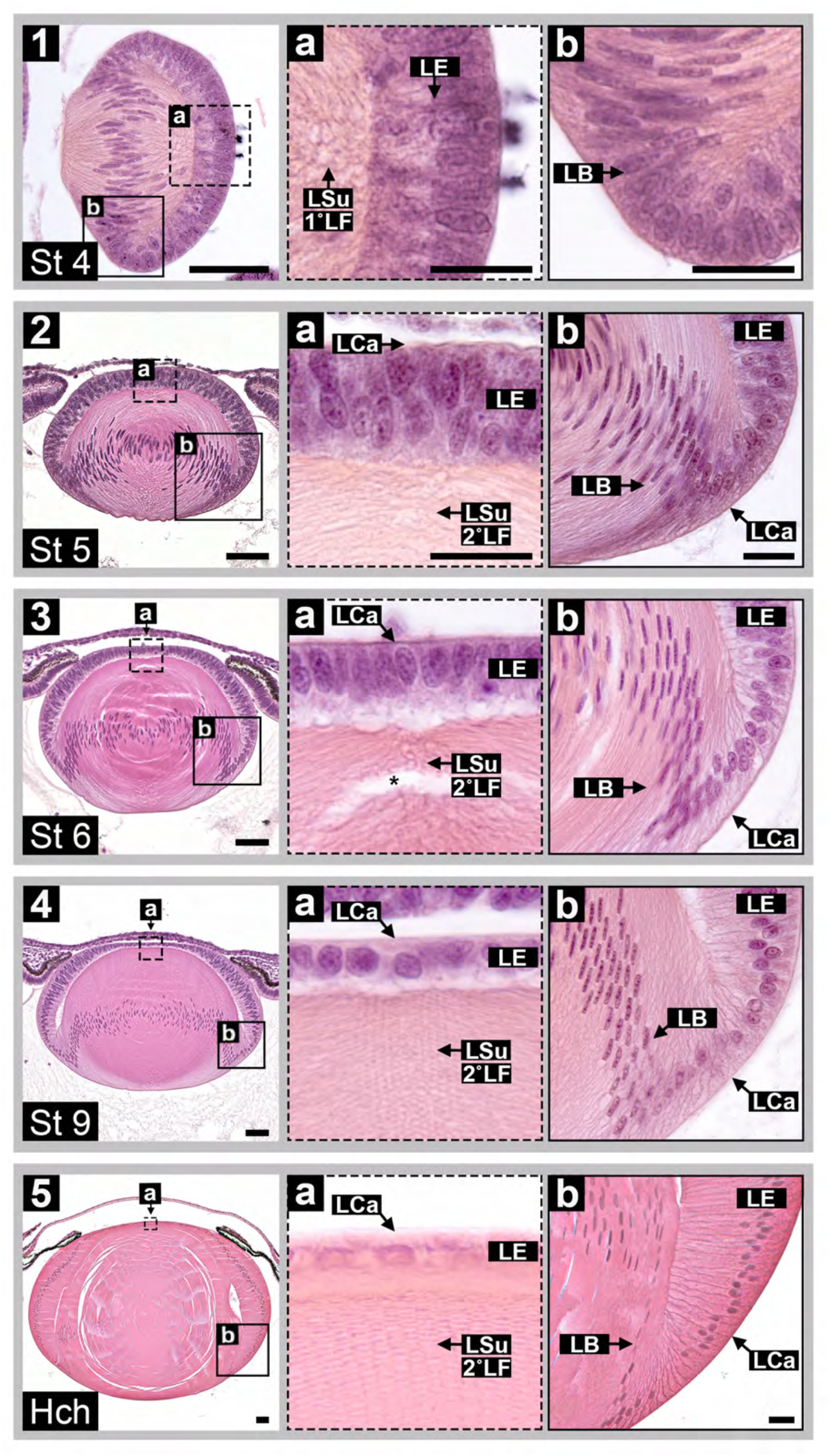
Lens maturation in the developing embryonic anole eye. (1-5) represent coronal (1) and horizontal (2-5) sections from the center lens from embryonic stages 4, 5, 6, 9, and hatchling; scale bars 100µm. High magnification views of lens epithelium (dashed boxes, 1.a-5.a) and lens bow (solid boxes,1.b-5.b) regions are shown; scale bars 20µm. Magnified images (a) and (b) from (2-5) with the exception of (5.b) are to scale with one another, respectively. Markers; LE – lens epithelium, LSu – lens suture, 1°LF – primary lens fiber, 2°LF – secondary lens fiber, LB – lens bow, LCa – lens capsule, and * indicates sectioning artifact.

After egg lay, the lens begins to increase in size with mitotic figures located in the epithelium (Figure 4, Figure 5). By St 5, secondary lens fibers are present at the lens equator and well-defined anterior and posterior sutures are evident (Figure 4.2,b). Within the anterior lens epithelium, which is approximately 2-4 cells thick (Figure 4.2,a), mitotic cells are observed throughout the epithelial layer anterior to the lens equator (Figure 5a). We did not observe any mitotic activity posterior to the lens bow, which suggests that these cells are undergoing differentiation and likely express crystallins. By stage 6, the anterior epithelial layer is thinner at its center and wider at its periphery (Figure 4.3, 4.3a). Proliferating cells are present throughout the central and peripheral lens epithelium as well as in the annular region. By St 9, when the anterior central epithelium becomes a single cell layer (Figure 4.4,a), cell division appears to be limited to the regions where the epithelium is still pseudostratified (Figure 5; diagramed in Figure 2). By St 11, elongating fiber cells within the lens core are undergoing denuclearization as pyknotic cells are evident within the lens nucleus (Figure 6.1,a); this process appears to be mostly complete by St 14 (Figure 6.2-3). An organelle-free zone (OFZ) is present in the lenses of St 15 embryos, which later expands in diameter as the lens continues to grow (St 16-18, Hch; Figure 6.4-5). In adults, presumptive lens fiber cells, identified by the presence of nuclei, are located throughout the center, peripheral, and annular pad epithelial regions of the lens (Figure 1.1,b,c). Cells undergoing denuclearization are also evident (Figure 1.b). These results suggest that the lens continues to grow throughout the life of the animal.

**Figure 5.**
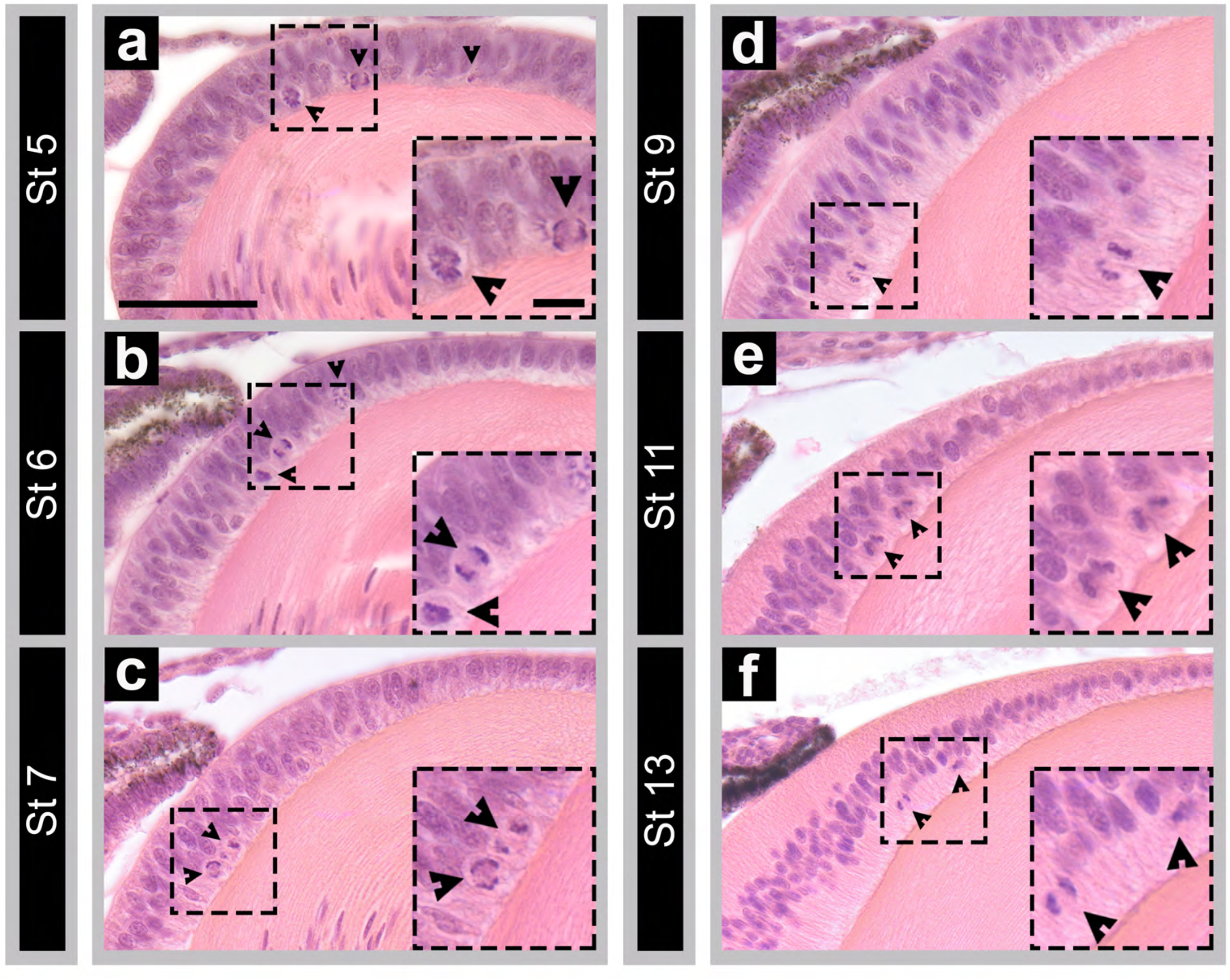
Cell proliferation in the anole lens epithelium. (a-f) shows horizontal sections from stages 5, 6, 7, 9, 11, and 13 embryos; scale bar 100 µm. High magnification inserts show miotic figures (arrow heads); scale bar 20 µm. Sections (a-f) and high magnification inserts are to scale with one another, respectively.

**Figure 6.**
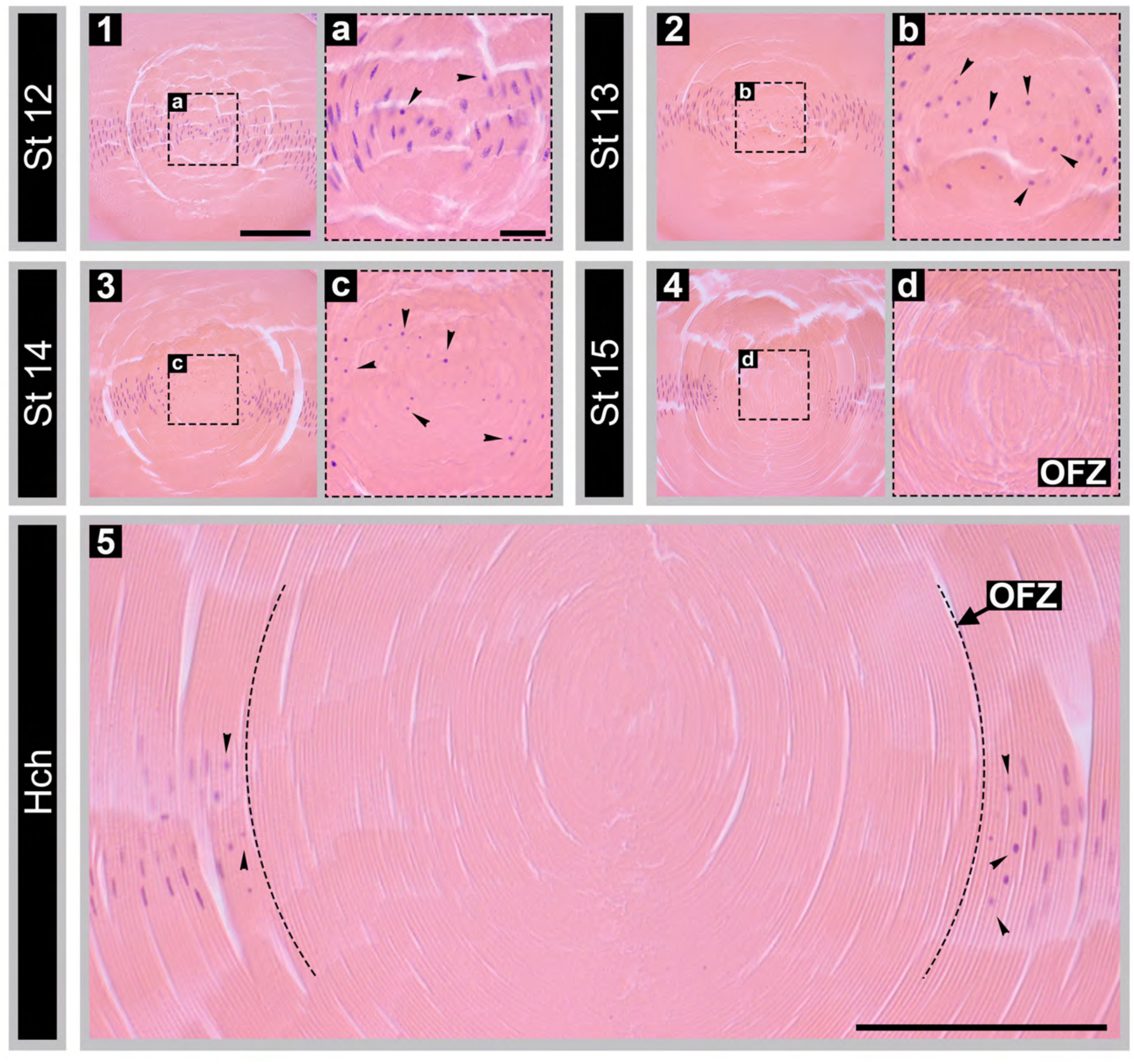
Denuclearization of the lens center. (1-5) depict horizontal sections of the lens nucleus; scale bar 100 µm. High magnification inserts show pyknotic figures (notches) in the lens center (OFZ – organelle free zone); scale bar 20 µm. Sections (1-4) and high magnification inserts (a-d) are to scale with one another, respectively.

### Cornea development

The surface ectoderm that will form the epithelial layer of the cornea is evident in embryos at PL 5 (Figure 3d). A few days later, at St 5, mesenchymal cells (presumptive migrating neural crest cells) are found underlying the corneal epithelium near the edge of the optic cup (Figure 7.1b). By St 6, the cornea has three distinct layers (Figure 7.2a). A thin acellular region, where Bowman’s membrane will eventually develop, separates the epithelial and endothelial layers and marks the beginning of primary corneal stroma formation (Figure 7.2a). This acellular region, however, is not long lasting. It disappears shortly before stage 9 as the first signs of keratocytes are detected (Figure 7. 3a). A analogous acellular zone has also been described for the developing cornea in chicks^19,28,29^, mice^30^ and primates^21,31^.

**Figure 7.**
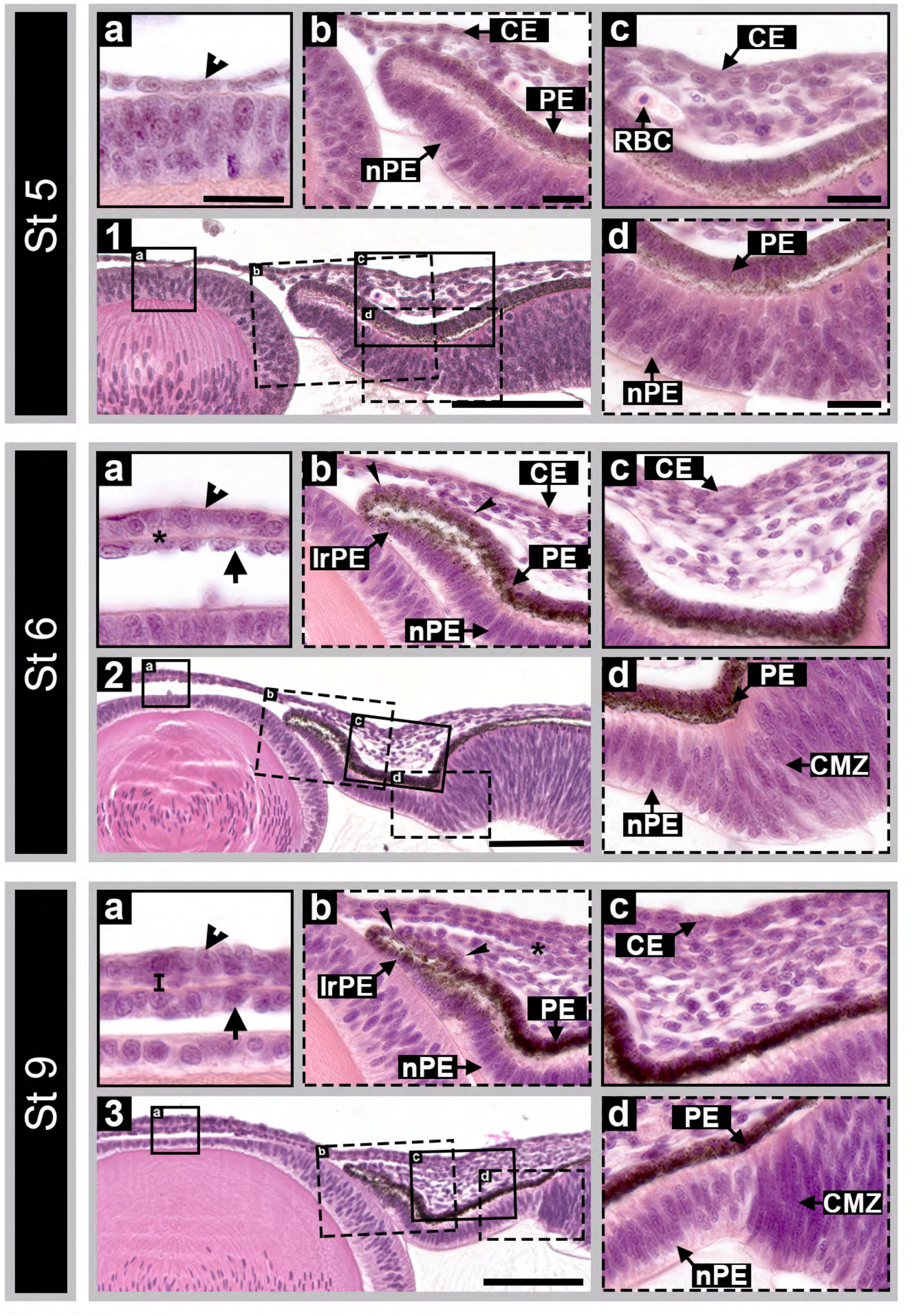
Early embryonic development of the anterior segment of the anole eye. (1-3) depict horizontal sections of the center eye from embryonic stages 5, 6, and 9; scale bars 100µm. High magnification inserts show cornea (a), iris (b), sclera sulcus (c), and ciliary body (d) regions of the developing eye; scale bars 20µm (a-d). Magnified inserts (1-3) are to scale with one another, respectively. Markers represent; CE – cornea epithelium, PE – pigment epithelium, nPE – nonpigmented epithelium, RBC – nucleated red blood cell, IrPE – iris pigmented epithelium, CMZ – ciliary marginal zone, arrowheads – cornea epithelium, * or bar – cornea stroma (2.a-3.a), arrow – cornea endothelium, * – iridocorneal angle (3.b), narrow notches – iris stroma anlagen.

Stromal expansion occurs between St 10-14. The number of presumptive keratocytes in the stromal layer increases to 3-4 cells deep between St 10-11 (Figure 8.4a) followed by deposition of collagen fibers (St 12-14). By St 14, the cornea reaches its maximal cellularity and width (Figure 8.6a). The stroma begins to condense during St 15, eventually shrinking nearly half its width by St 17 (Figure 9.8a, Figure 10). We postulate that the acellular layer immediately adjacent to the epithelium, is Bowman’s membrane. We could not however, readily identify a Descemet’s membrane. It is possible this layer is present but too thin to distinguish from the rest of the stroma in our histological sections. Accompanying these dynamic changes in stroma width are flattening of all the cornea cell layers, particularly the endothelial cells (Figure 10). By the time of hatching, the cornea is similar in all aspects of morphology and thickness to the adult.

**Figure 8.**
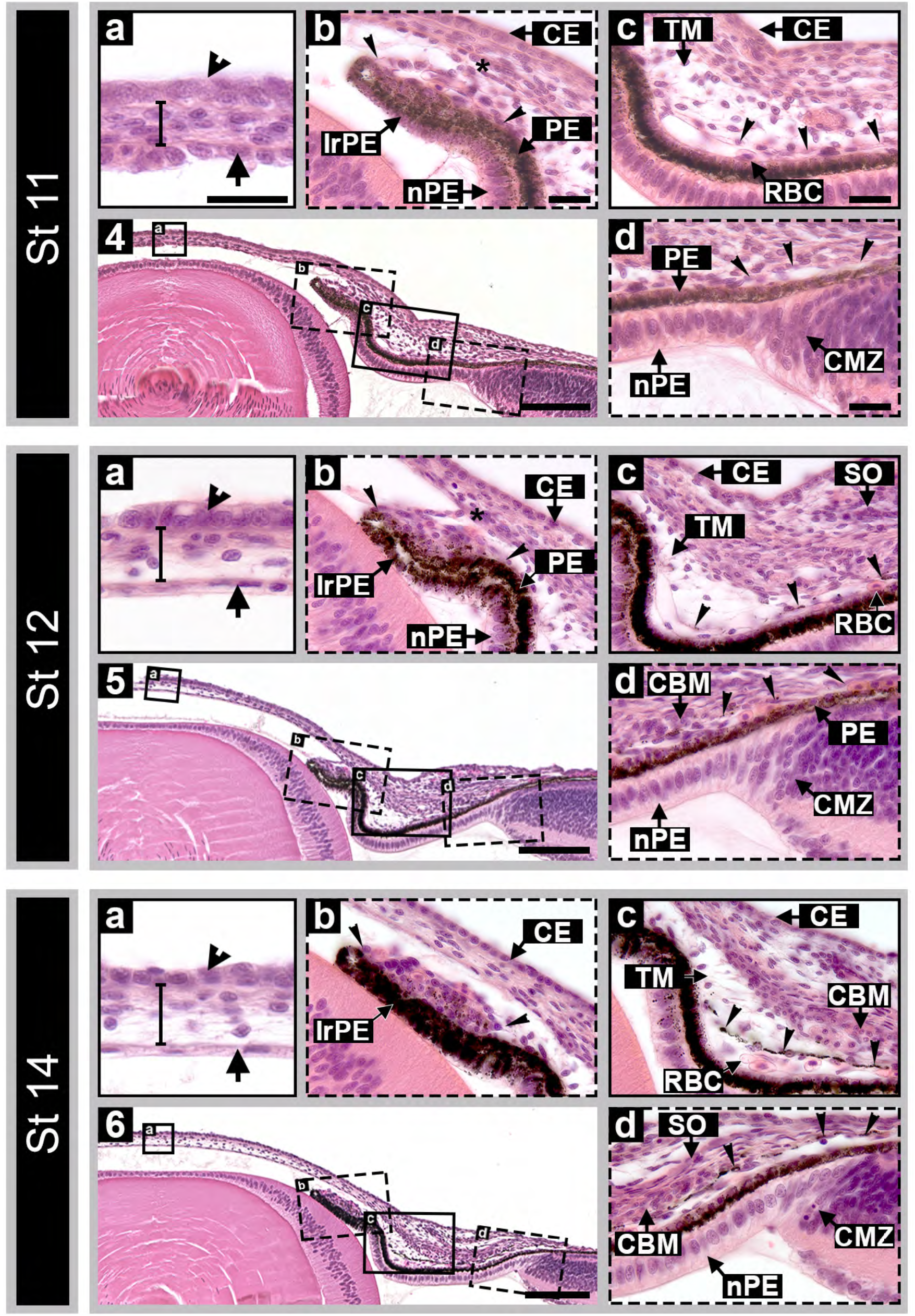
Mid embryonic development of the anterior segment of the anole eye. (4-6) depict horizontal sections of the center eye from embryonic stages 11, 12, and 14; scale bars 100µm. High magnification inserts show cornea (a), iris (b), sclera sulcus (c), and ciliary body (d) regions of the developing eye; scale bars 20µm. Magnified inserts (4-6) are to scale with one another, respectively. Markers; CE – cornea epithelium, PE – pigment epithelium, IrPE – iris pigment epithelium, nPE – nonpigmented epithelium, TM – trabecular meshwork, RBC – nucleated red blood cell, CMZ – ciliary marginal zone, SO – sclera ossicles, CBM – ciliary body muscle, arrowheads – cornea epithelium, bar – cornea stroma, arrow – cornea endothelium, * – iridocorneal angle (4.b-5.b), and narrow notches – iris or ciliary stroma anlagen.

**Figure 9.**
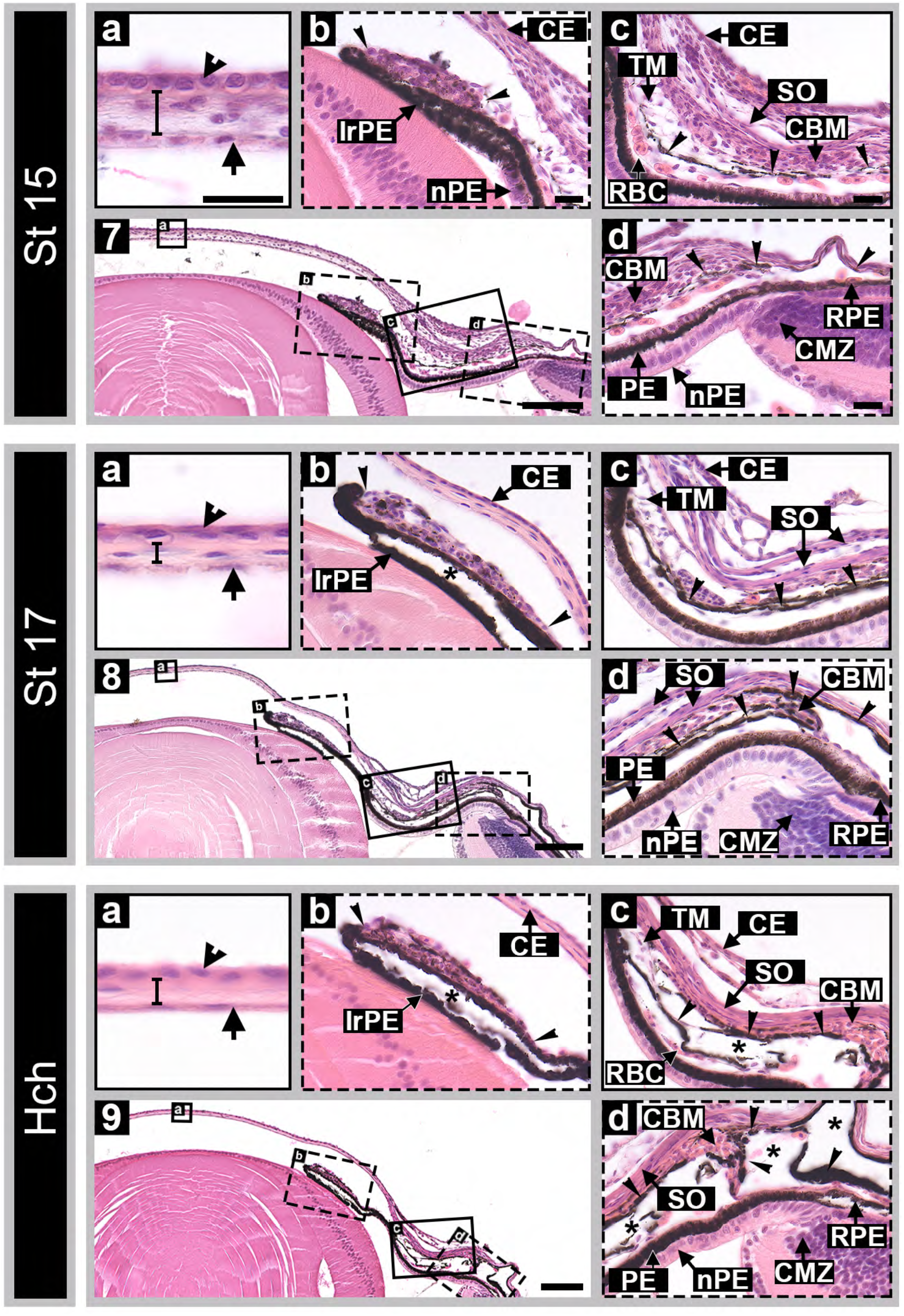
Late embryonic development of the anterior segment of the anole eye. (7-9) depict horizontal sections of the center eye from embryonic stages 15, 17, and hatchling; scale bars 100µm. High magnification inserts show cornea (a), iris (b), sclera sulcus (c), and ciliary body (d) regions of the developing eye; scale bars 20µm. Magnified inserts (4-6) are to scale with one another, respectively. Markers; CE – cornea epithelium, IrPE – iris pigment epithelium, nPE – nonpigmented epithelium, TM – trabecular meshwork, RBC – nucleated red blood cell, CBM – ciliary body muscle, SO – sclera ossicles, CMZ – ciliary marginal zone, RPE – retinal pigmented epithelium, arrowheads – cornea epithelium, bar – cornea stroma, arrow – cornea endothelium, narrow notches – iris or ciliary stroma anlagen, and * – sectioning artifacts.

**Figure 10.**
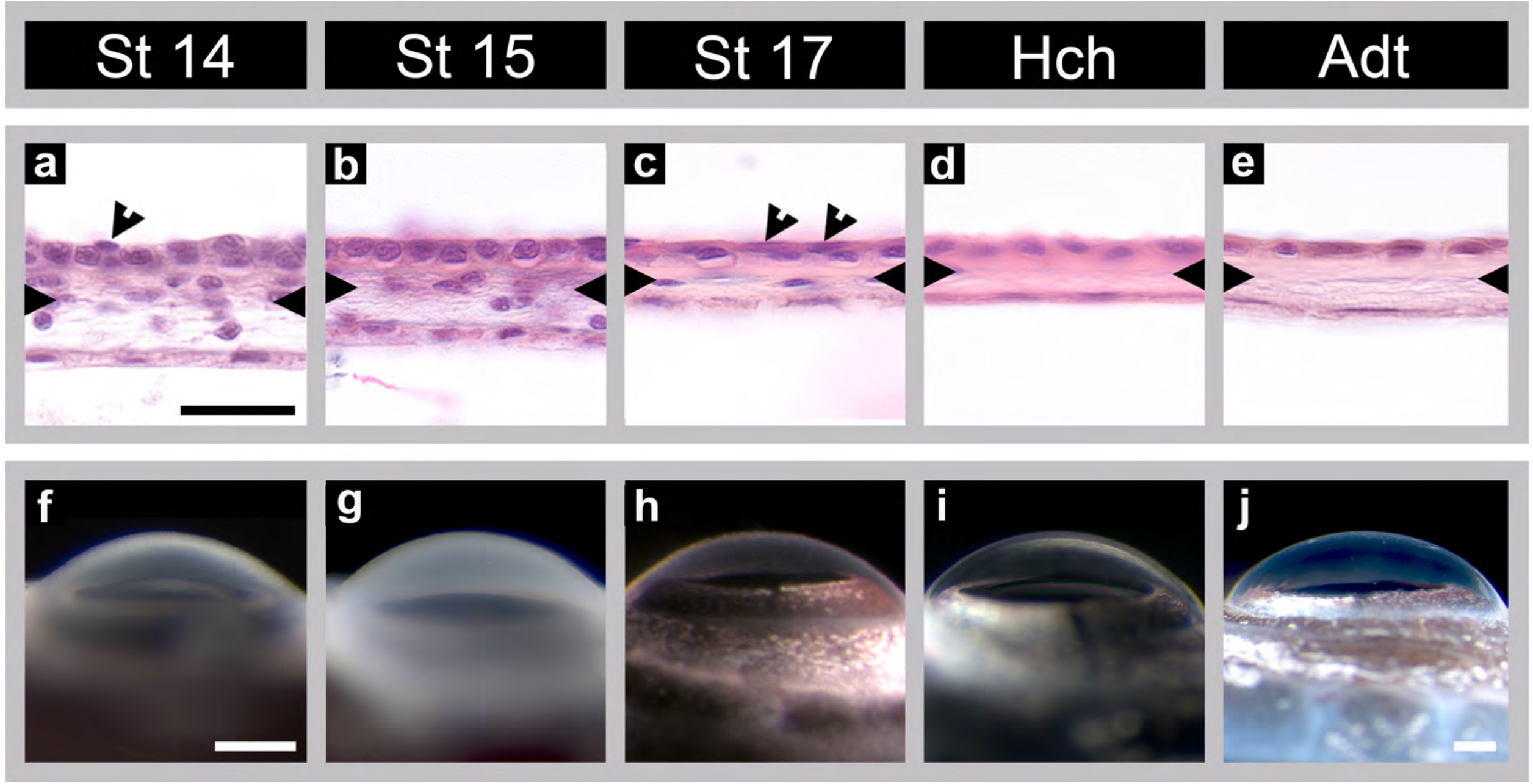
Corneal transparency increases as stroma condenses. Top panel (a-e) shows horizontal sections of the cornea center; scale bar 20 µm. Bottom panel (f-j) depicts the dorsal view from the same eyes shown in the panel above; scale bars 250 µm. Sections (a-e) and dorsal view images (f-i) are to scale with one another, respectively. Markers; arrowheads – 2 corneal epithelial cells on top of each other and notches – coalescent of bowman’s membrane boarder.

### Iris development

In St 5 embryos, the developing iris is distinguishable from the neural retina as an anterior marginal tapering of the optic cup (Figure 7.1b). Iris development initiates on the temporal side of the optic cup. By St 6, pigmentation can be seen accumulating along the developing outer pigmented epithelial cell layer of the presumptive iris and ciliary body. At this time, the outer pigmented epithelium is composed of a simple layer of short columnar cells, while the inner epithelium is pseudostratified and columnar (Fig 7.2b). By St 7-9, the iris primordium is morphologically distinct from the ciliary body (Figure 7.2b-3b). During this time, inner epithelial cells of the iris are also accumulating pigment, and by St 13-14, both iris outer and inner epithelial cell layers are heavily pigmented (Figure 8.6b). As the embryo nears hatching, the iris outer and inner epithelium becomes squamous-like in appearance (St 14-16) and by stage 17, greatly resemblances that of the hatchling (Figure 8.6b, Figure 9.7-9b).

The iris stroma is evident in St 6 embryos as mesenchymal-like cells that are closely associated with the iris outer pigmented epithelium (Figure 7.2b). By St 9 the cells appear as a simple cuboidal cell layer (Figure 7.3b). Cells adjacent to the outer pigmented epithelium are smaller and have round nuclei. Presumptive muscle fibers in the iris stroma are evident by St 13, and exhibit mature characteristics by St 17-18 (Figure 9.8b).

### Ciliary body development

The initial development of the ciliary body and iris occur concordantly. The ciliary body proper is morphologically evident on the temporal side of the optic cup in St 5 embryos and on the nasal side by St 6 (Figure 7.2, 4.2d, Figure 11.1a, 2b). The start of the ciliary body on the retina side is at the abrupt bend in the epithelium at the rim of the optic cup and extends distally to the second convex flexure, which is adjacent to the lens (Figure 7.2, 7.2c, 7.2d). The region distal to the second flexure is the iris. Shortly after the bend in the epithelium appears at the distal edge of the optic cup, the cells of the inner non-pigmented epithelium begin to organize into a simple cell layer, delineating the neural retina from the ciliary body (Figure 7.1d-3d, Figure 11.3c-4b). By St 16, the epithelium has become cuboidal in appearance (Figure 9.7d-8d). Pigmentation of the presumptive ciliary body is first observed at St 4 in the temporal region of the eye and later in the outer epithelial cells present in the sclera sulcus (St 5) (Figure 7.1c,d). Unlike the iris, the inner epithelial layer of the ciliary body is mostly devoid of pigmentation; however, cells near the iris-ciliary boundary have some pigment granules.

**Figure 11.**
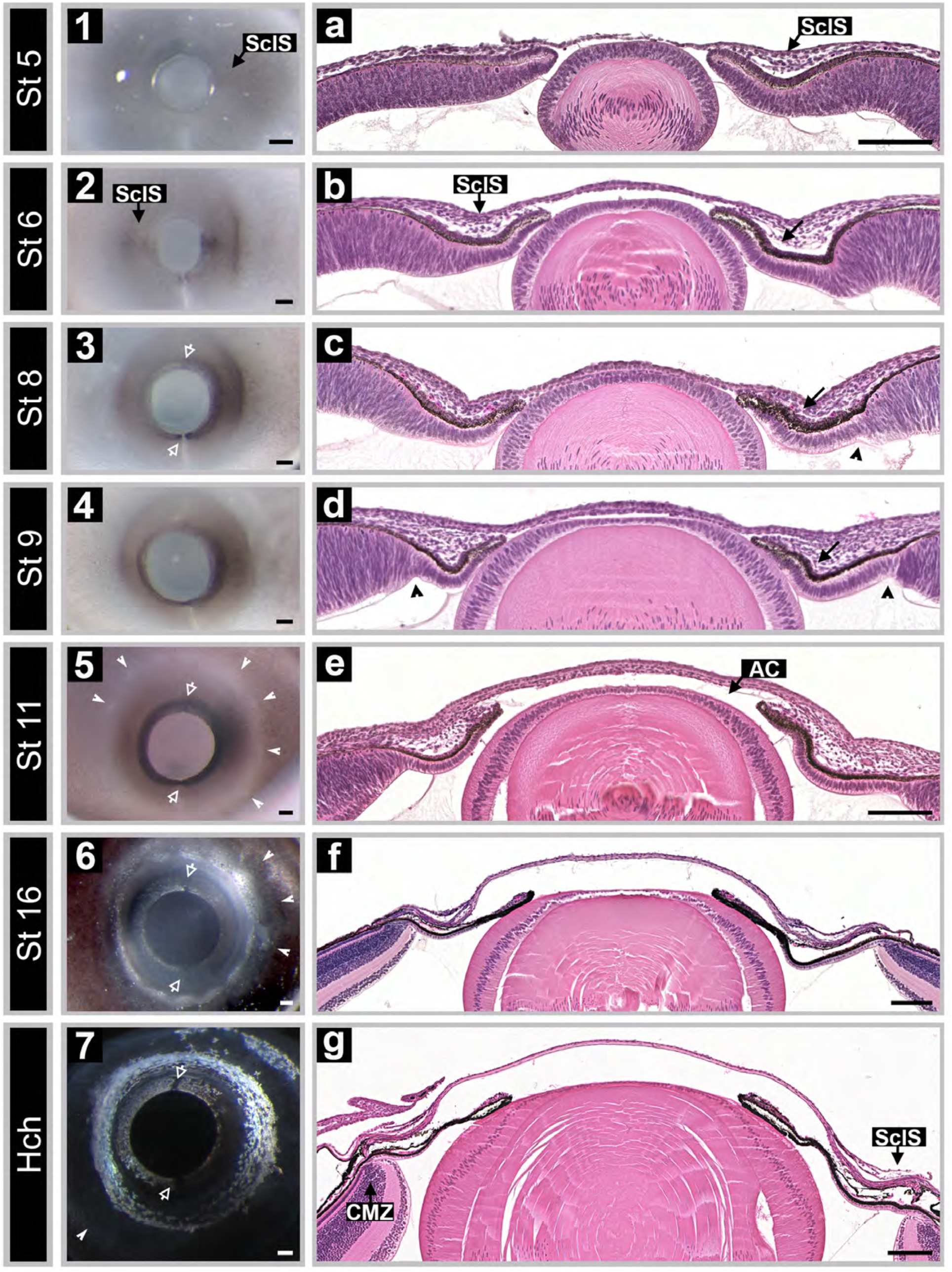
Spatiotemporal patterning of the anterior segment. (1-7) shows a lateral view of the left eye from embryonic stages 5, 6, 8, 9, 11, and 16 as well as the hatchling; scale bars 100 µm. (a-g) depicts horizonal sections of the anterior segment; scale bar 100 µm. Sections (a-d) are to scale with one another. Markers; SclS – sclera sulcus, AC – anterior chamber, arrows – iris flexure, arrow heads – ciliary margin zone, notches (white) – conjunctive papilla or sclera ossicles, and open arrows (white) – dorsal and ventral iris notches.

### Development of Iris and Ciliary body

The foremost observation we made while investigating the development of the iris and ciliary body relates to their relative size and spatiotemporal patterning. We observed that the temporal areas were much longer than the nasal regions and that the temporal side undergoes remodeling and differentiation prior to the nasal side (Figure 11). This patterning was evident shortly after anterior marginal tapering of the optic cup. A prominent convex flexure, known in lizards as the sclera sulcus, forms along the anterior margin and is first detected in the temporal region at stage 5 and later in the nasal region by stage 6 (Figure 11.1a-2b). This trend continues to hold at later timepoints when the iris and ciliary epithelium reorganize, forming the ciliary marginal region and undergoing pigmentation (Figure 11.3c-4d). For the most part, the development of nasal region appears to lag behind the temporal area by 1-2 stages (Figure 11). We observed a similar spatiotemporal pattern in the developing anole neural retina (*Rasys* et al., *in preparation*).

### Embryonic Sclera Ossicles & Ciliary muscle

Scleral ossicles are neural crest-derived membrane bones. The start of sclera ossicle formation can be seen in embryos as early as St. 11 as a pale ring of papilla in the conjunctiva surrounding the cornea and iris (Figure 11.5-6; see also, *Anolis sagrei* eye development poster in Supplementary Data by Rasys et al.^18^). In cross-section, condensing mesenchymal cells can be visualized and are located just posterior to the cornea-conjunctiva epithelium (Figure 8.4c). Between St 12-13, these cells organize into flat sheets approximately a few cells thick, extending initially from the sclera proper to the iridocorneal angle in an overlapping radial pattern (Figure 8.5c, 6c). By stages 14-15, the sclera ossicle anlagen begins to take on a cartilaginous-like appearance (Figure 8.6c, Figure 9.7-8c,d) and by stage 17, lacuna can be observed around cell nuclei in histological sections. Prior to this event, sclera extension in the temporal region takes place (St 10-14) and by stages 16-17, temporal asymmetry is easily seen.^18^ As the embryo continues to develop prior to hatching, the sclera ossicles thicken and expand from the sclera proper to the corneal boundary (Figure 9.8-9c,d).

In concert with sclera ossicle development, mesenchymal cells that give rise to the presumptive ciliary muscles are located adjacent to the developing ossicle (Figure 7.3c, Figure 8.4-5c,d). At stage 12 these cells occupy a broad region extending the entire length of the sclera sulcus. Ciliary muscle differentiation becomes evident between stages 13-14 as a condensation of the mesenchyme. Muscle fibers are present by stage 15. By stage 17, ciliary muscle resembles that of the hatchling (Figure 9.7-9c,d). Zonular fibers are detected at the time of hatching.

### Development of the Trabecular Meshwork

The iridocorneal angle is clearly evident in embryos at St. 9 as a separation of the iris stroma from the cornea, forming the anterior chamber (Figure 7.3, Fig 11.4). By St 11, a knot of mesenchymal cells separates the anterior chamber from a loose arrangement of cells sandwiched between the ciliary body and limbus (Figure 8.4c). We are defining this loose arrangement of cells as the trabecular meshwork in lizards, which bears resemblance to that of the early chick. ^26^ By stage 16-17, the trabecular meshwork appears to be largely consolidated near the iris-ciliary boundary and resembles that of the adult (Figure 9.7-8c). Bordering the ciliary muscle and adjacent to the trabecular meshwork are pigmented vascular endothelial cells that extend from the iridocorneal angle into the choroidal layer (Figure 8.6c,d, Figure 9.7-9c-d), which may serve to drain the aqueous humor.

## Discussion

Here, for the first-time, we describe in detail anterior eye development in a reptile, *Anolis sagrei*. We began our study by performing a detailed histological analysis of the structures of the adult eye. Conceptually, this starting point permits structural comparisons of the anterior segment in the brown anole with other species. A catalog of morphological similarities and differences between anole species and, more broadly, other reptiles can be used to index how representative the brown anole is as a reptile model for eye development. Pragmatically, this foundational work is required to support the study of anterior eye development in reptiles. Although there are a number of published reports on the reptilian eye, most are in the older literature and deal primarily with external anatomy and/or the structure of the adult retina.^1,9^ As a consequence, there is a paucity of detailed comparative information about the anterior eye in reptiles and its development. The data that does exist is mostly in the form of summary sketches that capture nicely the key morphological features of the anterior eye but lack the cellular detail useful for contemporary developmental studies.

Comparative assessments can be made at the gross anatomical level. *A. sagrei* possesses features found in lizards and turtles, but not snakes, amphibians or placental mammals.^1,9^ These include the presence of scleral bone and cartilage, a lens with an annular pad, and skeletal rather than smooth muscles in the iris and ciliary body. These unique features are also found in birds.^1^ Compared with other anoles, we find that the anterior segment of *A. sagrei* is structurally similar to that in *A. carolinensis* (data not shown) and *A. lineatopus* ^9^.

The most striking difference in the anterior segment of anoles compared to most other reptiles, birds and mammals is the structure of the adult cornea. In *A. sagrei*, the corneal epithelium is largely composed of a single basal cell sheet (Figure 1.1a). This was surprising as the corneal epithelium in many vertebrates is a stratified squamous epithelium consisting of several layers of cells. For example, the adult epithelium in chickens is typically composed of 6-7 cell layers,^19^ while mice have 4-5 cell layers,^20^ and humans have 5-7 layers.^21,32^ Among the reptiles that have been examined, the number of cell layers varies depending on the species. For example, the cornea epithelium is 2-3 cells thick in grass snakes (*Natrix natrix*) and sand lizards (*Lacerta agilis*), and 3-4 cell layers thick in pond tortoises (*Emys orbicularis*).^1,22^ The anole corneal stroma is also thinner relative to that in other reptiles, birds, and placental mammals.^19-21,28,30-37^

The physiological impact of a thin cornea, and especially the epithelial layer, is unclear. The cornea acts as an external barrier to protect the eye and is a critical element in establishing the overall optical properties of the eye. In addition to its barrier function, the epithelial layer plays a critical role in maintaining the clarity of the cornea. The epithelial layer is constantly turning over. In mammals, a population of stem cells, located at corneal-scleral junction, give rise to the corneal epithelial cells that maintain the ocular surface. Newly born epithelial cells migrate radially into the basal cell layer of the epithelium. As cells desquamate from the corneal surface, cells from deeper layers migrate towards the surface. As cells migrate to more superficial layers, they also undergo changes in their morphology and physiological properties. In birds and mammals, basal cells that leave the basement membrane normally take on a polyhedral shape and begin expressing keratins. These cells then take on a squamous morphology, with flattened nuclei, as they move towards the corneal surface. It will be interesting to determine the mechanisms that maintain the ocular surface in anoles.

The development of the anterior segment in *A. sagrei* progresses in a similar fashion to that observed for all vertebrates. The first structure to form is the lens, followed by the cornea and later the iris, ciliary body, and structures responsible for draining the aqueous humor from the eye.^20,30,37-42^. However, the ways in which the structures of the anterior segment develop are more similar to the chicken^19,28,33-36^ than mice^20,30^ or haplorhini primates.^21,31,37^. This correlation is consistent with the phylogenetic relationship of reptiles to birds and mammals.

Based on our histological analysis it is unclear if lizards possess Schlemm’s canal. In his landmark comparative work on the vertebrate eye, Walls (1942) stated that Schlemm’s canal was located at the iridocorneal angle of the lizard eye as it is in mammals. In humans and other primates, Schlemm’s canal is an endothelium-lined vascular structure immediately adjacent to the juxtacanalicular region of the trabecular meshwork and encircles the cornea.^25,42^ The endothelial cells that make up the inner and outer walls of Schlemm’s canal have distinctive characteristics that can be seen in histological sections. Using these characteristics as a guide, we have not been able to identify a comparable structure in either the adult or developing eye of *A. sagrei*. Our failure to identify Schlemm’s canal in the brown anole is consistent with the observations reported for *A. lineatopus* by Underwood (1970), in which he noted that Schlemm’s canal “is poorly defined in Anolis.”

The primary functions of Schlemm’s canal are to maintain intraocular pressure and serve as a conduit for the aqueous humor to flow into the venous system.^25,42^ We propose that this function is served in anoles by the vasculature from the choroid region that extends between the ciliary body and the ciliary muscles to the iridocorneal angle. Elucidating the cellular, molecular, and genetic mechanisms that regulate the development of the aqueous humor outflow pathways in *anolis* lizards may provide insights into the different ways this important fluid handling problem can solved in nature.

